# APOBEC3F is the main source of editing identified during the 2022 outbreak of human monkeypox virus

**DOI:** 10.1101/2023.01.06.522979

**Authors:** Rodolphe Suspène, Kyle A. Raymond, Laetitia Boutin, Sophie Guillier, Frédéric Lemoine, Olivier Ferraris, Jean-Nicolas Tournier, Frédéric Iseni, Etienne Simon-Lorière, Jean-Pierre Vartanian

## Abstract

On May 6, 2022, a powerful outbreak of monkeypox virus (MPXV) had been reported outside of Africa, with many continuing new cases being reported around the world. Analysis of mutations among the two different lineages present in the 2021 and 2022 outbreaks revealed the presence of G->A mutations occurring in the 5’GpA context, indicative of APOBEC3 cytosine deaminase activity. By using a sensitive PCR (3D-PCR) method allowing differential amplification of AT-rich DNA, we demonstrate that G->A hypermutated MPXV genomes can be recovered experimentally from APOBEC3 transfection followed by MPXV infection. Here, among the 7 human APOBEC3 cytidine deaminases (A3A-A3C, A3DE, A3F-A3H), only APOBEC3F was capable of extensively deaminating cytidine residues in MPXV genomes. Hyperedited genomes were also recovered in ~42% of analyzed patients, indicating that editing is part of the natural cycle of MPXV infection. Moreover, we demonstrate that substantial repair of these mutations occurs. Upon selection, corrected G->A mutations escaping drift loss contribute to the MPXV evolution observed in the current epidemics. Thus, stochastic or transient overexpression of *APOBEC3F* gene exposes the MPXV genome to a broad spectrum of mutations that may be modeling the mutational landscape after multiple cycles of viral replication.

---

Monkeypox is a neglected, zoonotic infectious disease caused by the monkeypox virus (MPXV). This virus belonging to the *Orthopoxvirus* genus consists of a double-stranded, 197-kb DNA genome. MPXV was first isolated in 1958 from a *Macaca fascicularis*, having originated from Singapore and imported to Copenhagen, which subsequently caused an outbreak in captive *Cynomolgus* monkeys (Pv et al., 1959). By 1970, the virus proved capable of a zoonotic jump to humans, but remained contained in Africa, causing isolated episodes of infection. This was until 2003, when an outbreak related to the import of small African mammals appeared in the United States ((CDC), 2003). In 2017, the largest monkeypox outbreak in West Africa occurred in Nigeria (Yinka-Ogunleye et al., 2019), after decades of no identified cases. From 2018 to 2021, several cases had been imported from Nigeria to non-endemic countries such as United Kingdom, Israel, and Singapore (Erez et al., 2019; Mauldin et al., 2022; Ng et al., 2019; Vaughan et al., 2018). In May 2022, a rapidly growing multi-country outbreak involving individuals without travel history to MPXV endemic countries was identified. By July 23^rd^ 2022, the WHO declared a Public Health Emergency of International Concern. As of December 19^st^ 2022, 82,999 MPVX confirmed cases have been reported (Centers for Disease Control and Prevention, 2022). International sequencing efforts revealed that the 2022 outbreak virus belongs to MPXV clade IIb (part of the formerly designated “West African” clade)(Happi et al., 2022) and forms a divergent lineage (B.1) that descends from genomes related to the 2017-18 outbreak in Nigeria (Figure 1) (Isidro et al., 2022).

**Figure 1.**
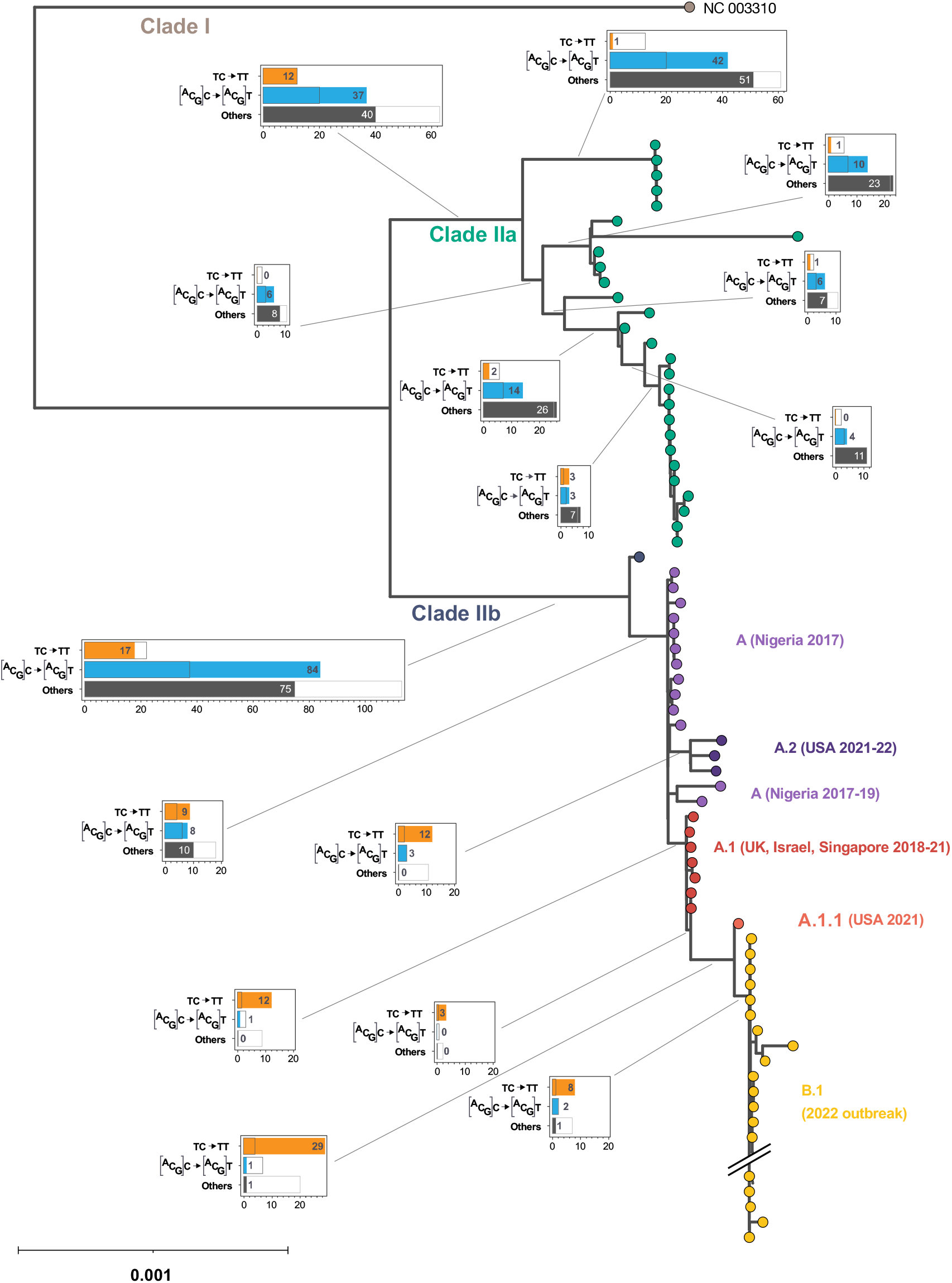
Phylogenetic analysis of substitutions within MPXV clade IIa and IIb. A Maximum likelihood tree was inferred using IQ-TREE2. The observed number of each type of mutations are indicated in the filled barplots along each branch of the phylogeny. The orange and blue boxes indicate respectively the C->T mutations occurring in the 5’TpC->TpT and 5’(A or C or G)pC->(A or C or G)pT contexts while the black boxes indicate all other types of mutations. Empty barplots represent the average number of mutations obtained over 100 simulations under the substitution model used for tree inference.

The extent of the divergence of the 2022 outbreak genomes from the related 2018-19 viruses is striking, considering the estimated substitution rate for Orthopoxviruses (Firth et al., 2010). The most accumulated mutations in the 2022 MPXV genomes (42 out of 47 mutations) correspond to cytidine (C) to thymine (T) (C->T) transitions, occurring in the 5’TpC context (5’TpC->TpT or 5’GpA->ApA in the reverse strand) (O’ Toole and Rambaut, 2022). This mutation signature is unique but pervasive in recent clade IIb genomes, with significant signal in sublineage branches (Gigante et al., 2022). Importantly, this signature is absent from clade I, IIa genomes or older clade IIb MPXV genomes (Figure 1) and a complete phylogeny with tip names is presented in Figure supplement 1. The tree displayed in Figure 1 presents the number and the nucleotide context of each type of mutations that occurred at different branches (between inferred ancestral sequences) and shows a clear signature starting from clade A.2 and A.1, that is not expected under the substitution model used for tree inference. These monotonous mutations, occurring in a specific context, reflects the footprint of an APOBEC3 cytidine deaminase (A3).

In the last two decades, it has emerged that the human A3 locus encodes six functional polynucleotide cytidine deaminases (A3A, A3B, A3C, A3F, A3G, A3H) (Jarmuz et al., 2002), originally described as innate cellular restriction factors against viruses and retroelements through the deamination of C to U in single-stranded DNA (ssDNA) (Harris et al., 2003; Lecossier et al., 2003; Mangeat et al., 2003; Mariani et al., 2003; Sheehy et al., 2002; Zhang et al., 2003). These deamination events occur preferentially in the context of 5’TpC, with the exception of A3G, which prefers 5’CpC dinucleotides (Beale et al., 2004; Bishop et al., 2004; Liddament et al., 2004; Suspène et al., 2004).

To determine whether A3 enzymes are capable of editing MPXV genomes, HeLa cells were transfected with A3A, A3B, A3C, A3F, A3G, A3H or with an empty plasmid (ep) used as a negative control, and then infected with a low passage 2022 MPXV isolate (Frenois-Veyrat et al., 2022).

Differential DNA denaturation polymerase chain reaction (3D-PCR) was used to detect and selectively amplify AT-rich edited genomes located in MPXV *B10R ER-localized apoptosis regulator* (*Cop-B9R*) gene (Suspène et al., 2005b). As shown in Figure 2a, 81.2°C was the minimal temperature that allowed amplification of the MPXV genome with A3A, A3B, A3C, A3G, A3H and ep. Specifically, it was possible with A3F overexpression to selectively amplify MPXV DNA at 80.7°C, indicating a higher mutational burden. The A3F enzyme was able to extensively deaminate MPXV DNA (Figure 2b), with the complete sequences presented in Figure supplement 2. Sequencing revealed extensive and monotonous cytidine deamination of both DNA strands (Figure 2c). We determined that the mean of G->A editing frequency was ~22% (range, 5-79% per clone). Analysis of the dinucleotide context of edited sites showed a strong 5’ effect favoring 5’GpA (5’TpC in the opposite DNA strand) typical of A3F enzyme (Beale *et al*., 2004; Bishop *et al*., 2004; Liddament *et al*., 2004; Suspène *et al*., 2004) (Figure 2d). By contrast, there was no pronounced 3’ nucleotide context, such as 5’CpG, thus ruling out a cytidine hypermethylation/deamination related phenomenon.

**Figure 2.**
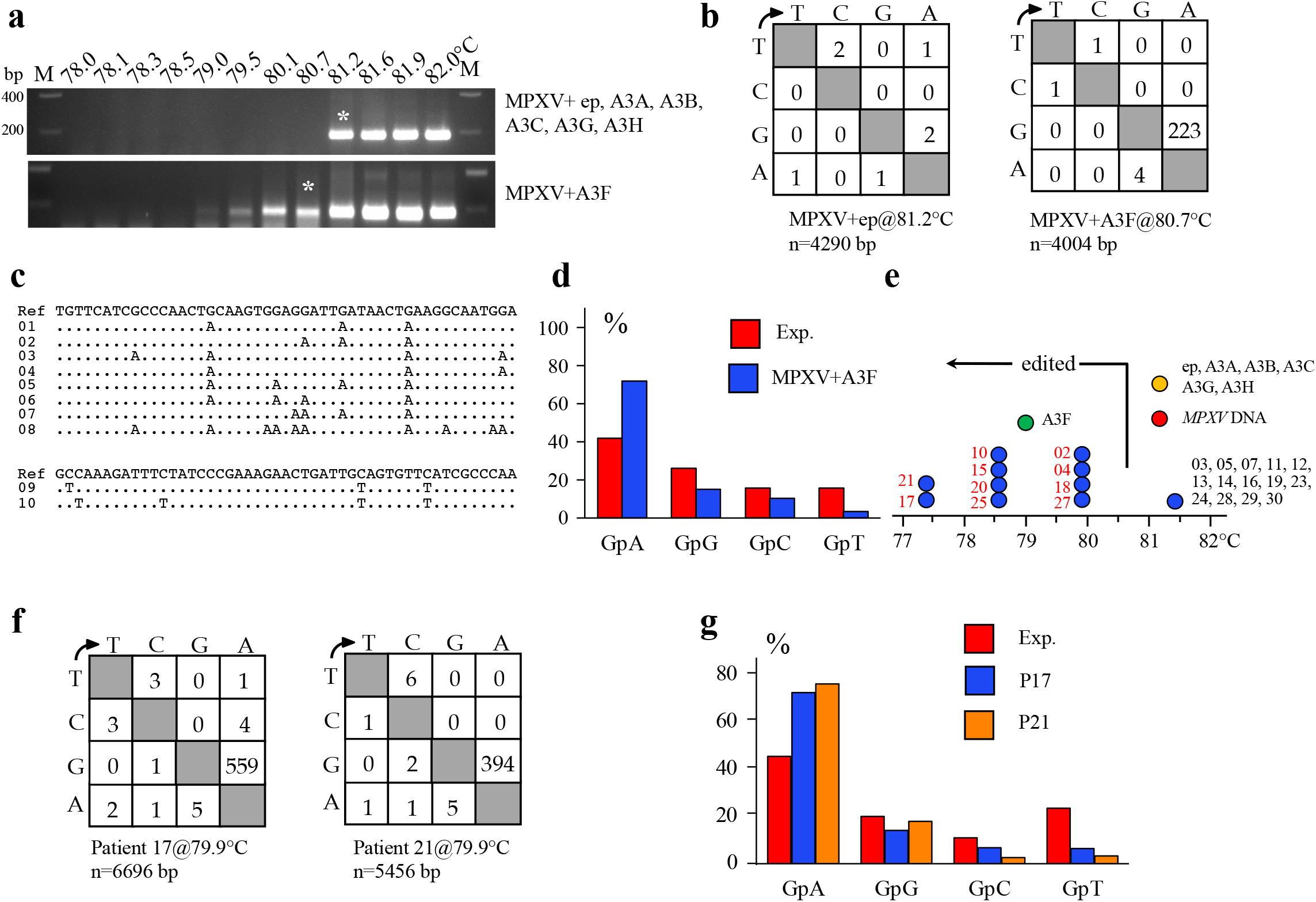
APOBEC3F editing of MPXV DNA. a) 3D-PCR recovered A3F- and ep, A3A-A3C, A3G, A3H-edited MPXV genomes down to 80.7°C and 81.2°C respectively. ep: empty plasmid vector and MPXV alone, M: molecular weight markers, bp: base pair. Asterisks refer to the samples cloned and sequenced. b) Mutation matrices for hyperedited *Cop-B9R* sequences derived from cloned 80.7°C and 81.2°C 3D-PCR products. n indicates the number of bases sequenced and corresponds to 28 and 30 sequences x143bp for ep or A3F+MPXV respectively. c) A selection of hypermutated A3F G->A and C->T edited MPXV genomes (Td = 80.7°C). Although editing may occur on both strands, the sequences are given with respect to the plus or coding strands. Only differences are shown. All sequences were unique, indicating that they corresponded to distinct molecular events. d) Bulk dinucleotide context of MPXV *Cop-B9R* gene fragment by A3F cytidine deaminase and compared to the expected values. The red bars represent the expected frequencies assuming that G->A transitions were independent of the dinucleotide context and correspond to the weighted mean dinucleotide composition of the reference sequence (accession MT903344). The blue bars represent the percentage of G->A transitions occurring within 5’GpN dinucleotides for the hypermutated sequences. e) Schematic representation of the denaturation temperature (Td) for the last positive 3D-PCR amplifications for 24 MPXV infected patients (dark blue circles). The arrow indicates the threshold Td (80.7°C) at which the samples are hypermutated. The samples in red were cloned and sequenced. The red, orange and green circles represent respectively the last Td of the MPXV DNA molecular clone corresponding to the reference sequence, ep, A3A-A3C, A3G, A3H and A3F. f) Mutation matrices for hyperedited *Cop-B9R* sequences derived from patients 17 and 21 at 79.9°C, n indicates the number of bases sequenced and corresponds to 27 and 22 G->A sequences x 248bp for patients 17 and 21 respectively. g) Bulk dinucleotide context of MPXV *Cop-B9R* gene fragment for patients 17 and 21and compared to the expected values.

Next, we tested whether edited MPXV genomes could be detected in patients. 3D-PCR was performed on DNA extracted from 24 MPXV-infected patients presented in Table supplement 1. When considering the most selective denaturation temperature (Td) by 3D-PCR, we observed that 10 of the 24 (~42%) patient samples analyzed exhibited hypermutated sequences. (Figure 2e, samples in red validated by sequencing, samples with a Td>80.7°C being non hyperedited).

In light of these data, we decided to analyze patients 17 and 21 in detail, as they presented the most hyperedited viruses, as visualized by the last Td obtained and the mutation matrices (Figure 2E, 2F). As observed in Figure 3a and Figure supplement 2, at Td=79.9°C, G->A and C->T hyperedited genomes were found in patients 17 and 21, proving that both DNA strands could be edited (Taq/79.9°C). The mean editing frequency of G->A was similar between the two patients ~38% and ~32% (range, 14-47% and 16-58% for patient 17 and 21 respectively). By analyzing the editing frequencies obtained with a Td= 95°C, we were able to determine that the *in vivo* frequency of hypermutants is ~1.75% (1 hypermutant detected among 58 sequences, patient 17), proving that this phenomenon is not a rare event (Figure 3a, Figure supplement 2).

**Figure 3.**
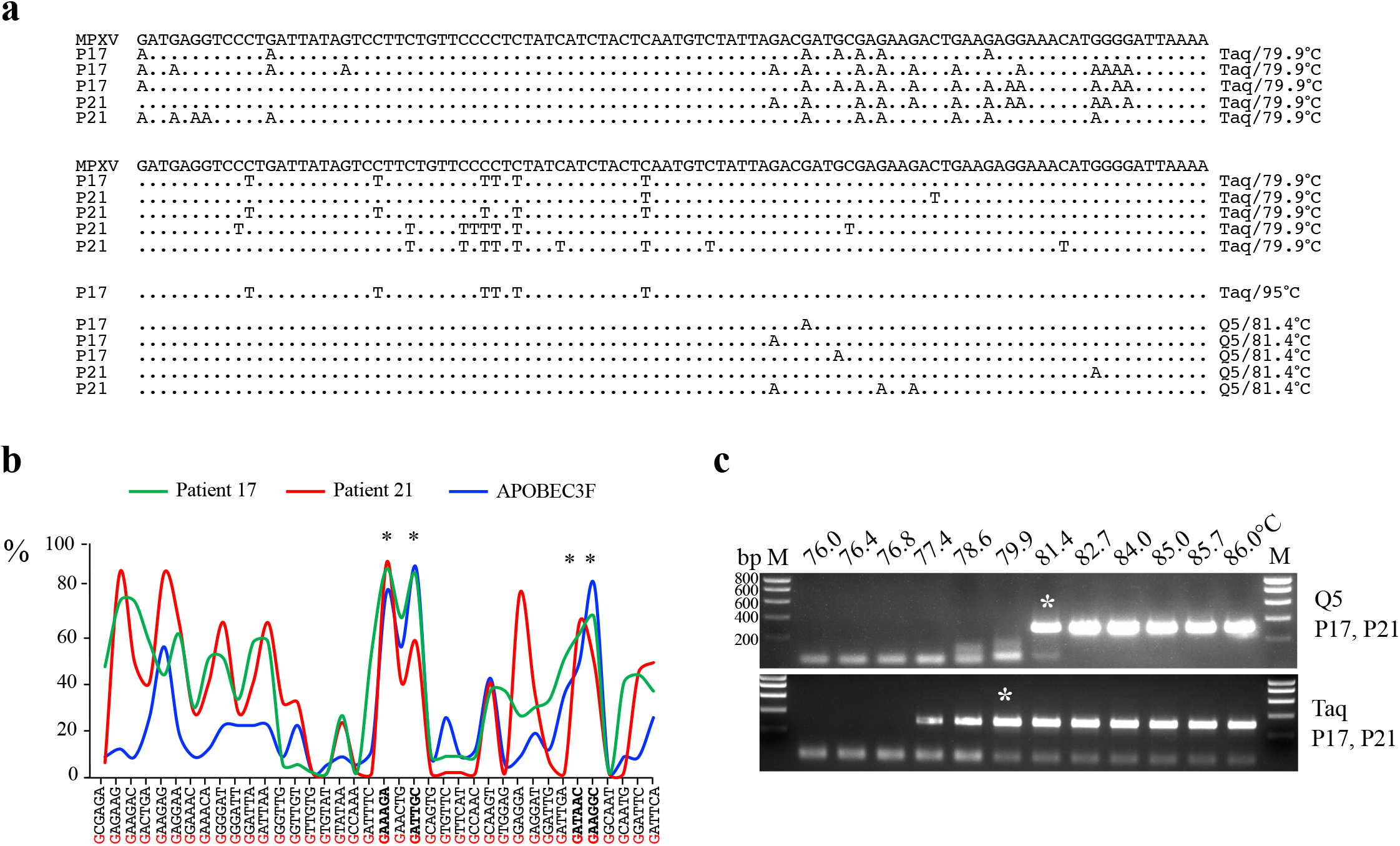
Hypermutation of MPXV DNA *in vitro* and *in vivo*. a) A selection of hypermutated MPXV sequences from patients 17 and 21 (Td = 95°C, 81.4°C and 79.9°C) is shown and compared with the reference. The sequence is given with respect to the viral plus strand. Only differences are shown. All sequences were unique, indicating that they corresponded to distinct molecular events. b) Site specific frequency analysis of editing from A3F transfected cells and from patients 17 and 21 recovered by 3D-PCR. The stars represent overlapping hot spot with the context in bold. Analysis was performed from nucleotide 168303-168445, accounting for 38 Gs. c) 3D-PCR amplification across a 76-86°C gradient using either Taq or Q5 DNA polymerase, the latter fails to amplify DNA containing dU, the product of A3 deamination. Asterisks indicate the PCR products cloned and sequenced, M: molecular weight markers, bp: base pair.

The dinucleotide context associated with G->A hyperedited genomes *in vivo* showed an overall preference for 5’GpA (5’TpC in the opposite DNA strand, Figure 2g), which is comparable to those detected *in vitro* (Figure 2d). By comparing the mutational frequency of MPXV DNA from patients 17 and 21 at each of the 38-target G residues with those derived from the A3F transfection, significant correlations with the presence of mutational hotspots were observed between the *in vitro* and *in vivo* edited sites (Figure 3b, Figure supplement 2). This collection of evidence clearly indicates that the main mutator enzyme of the MPXV genome *in vivo* is A3F, however, it is not excluded that other A3s could be involved in MPXV editing both *in vitro* and *in vivo*.

Upon A3-induced cytidine deamination, DNA bearing multiple dU residues might be corrected by the MPXV UNG pathway or, if copied on the opposite strand, the resulting dU:dA base pair might be proofread to dT:dA. To explore this issue, we performed the first round PCR using either Q5 or Taq polymerases on extracted DNA from patients 17 and 21 lesions. Like all archaeal DNA polymerases, Q5 is unable to amplify DNA templates bearing dU (Wardle et al., 2008), if only one of the many existing dUs is uncorrected, the genome will not be amplified by the Q5. To assess the efficacy of the MPXV UNG repair system, 3D-PCR products obtained at Td=79.9°C and 81.4°C with Taq and Q5 polymerases respectively were cloned and sequenced (Figure 3c, Figure supplement 2). As can be observed, Q5 amplification recovered less edited DNA than Taq polymerase, indicating that some dU:dA pairs can be repaired to dT:dA. These observations indicate that substantial repair does occur, demonstrating that upon selection, corrected G->A mutations that escape drift loss are contributing to the MPXV evolution as observed in the current epidemics.

We have demonstrated that the MPXV viral mutations observed during the successive outbreaks of 2017-19 and 2022 are in part the consequence of A3F cytidine deaminase activity. The library of hypermutated MPXV genomes identified presented a range of editing from ~5 to 79% (Figure supplement 2). These data show that there is a range of mutations among the analyzed sequences, demonstrating a balance between A3 associated hypo- and hyperediting of the MPXV genome. Hyperediting frequency detected can be phenomenally high, >10^−1^ per base, so much so that they result in defective viruses. On the contrary, hypoediting rates are lower, perhaps at the order of 10^−6^ per base. Obviously, only limited deamination of viral genomes is likely to generate viruses presenting fitness compatible with further transmission, and thus to have the chance to be fixed in a successful viral lineage.

A number of DNA viruses have been documented to exhibit mechanisms that limit or counteract the effects of A3 enzymes, such as coding for a viral interactor, preventing A3 incorporation into virions, replicating in cells with low or absent A3 expression, or replicating in preferred subcellular locations. The efficacy of these mechanisms may differ in MPXV lineage B viruses compared to other lineages, but it is also conceivable that ecological shifts, including sustained human-to-human transmission in sufficient numbers, may have led to the detectable occurrence of this process.

Further genomic surveillance will show whether fortuitous founder events contributed to the genetic make-up of the genomes detected in the 2022 outbreak, and whether this process will continue to play a role in the evolution of MPXV in the ongoing outbreak. Indeed, this protective mechanism may eventually contribute to evolutionary jumps or increased evolutionary rates, that could be associated with viral immune escape or resistance to treatment. Understanding the mechanisms involved behind this editing process could lead to promising new antiviral approaches against MPXV.

## Materials and Methods

### Cells and virus

HeLa cells (Human epithelial cells, ATCC CCL-2) were maintained in Dulbecco’s modified Eagle’s medium (DMEM, Gibco), supplemented with heat-inactivated fetal calf serum (10%), penicillin (50 U/mL), and streptomycin (50 mg/mL) and were grown in 75-cm^2^ cell culture flasks in a humidified atmosphere containing 5% CO_2_.

A virus from the 2022 MPXV outbreak was isolated, and a stock was produced on Vero cells (Frenois-Veyrat *et al*., 2022). Passage 2 stock was titered by plaque forming assay in 24-well plates and sequenced. The obtained titer was 2×10^6^ PFU/mL.

### Transfection, viral infections, 3D-PCR, cloning and sequencing

Transfections were performed independently in triplicate on HeLa cells. Briefly, 2.5×10^5^ HeLa cells were transfected with 1μg of individual plasmids encoding either A3A, A3B, A3C, A3F, A3G, A3H or an empty plasmid (ep) as control, using jetPRIME (Polyplus). At 24 hours post transfection, HeLa cells were infected with MPXV at MOI 1 for 24 hours. Total DNA from HeLa infected cells were extracted with the QiAmp DNA Mini kit (Qiagen). Differential DNA denaturation polymerase chain reaction (3D-PCR) was performed on an Eppendorf gradient Master S programmed to generate a 78-82°C or 76-86°C gradient in the denaturation temperature. This technique relies on the fact that AT-rich DNA denatures at a lower temperature than GC-rich DNA (Suspène *et al*., 2005b) and has proven successful at selectively amplifying APOBEC3-edited viral DNA for Hepatitis B virus (HBV) (Suspène et al., 2005a; Turelli et al., 2004), papillomaviruses (HPV) (Vartanian et al., 2008), herpesviruses (HSV, EBV) (Suspène et al., 2011) as well as for viruses with a DNA intermediate, such as retroviruses (Delebecque et al., 2006; Mahieux et al., 2005).

A fragment of *B10R ER-localized apoptosis regulator* (*Cop-B9R*) gene position (nucleotide 168241-168906, accession MT903344), was amplified by employing a nested procedure. For the *in vitro* experiments, the first-round primers were 5’ GACTAAATTTCTCGGTAGCACATCGAA and 5’ GGGACACCTGTATTCATGTTACTGAA. First PCR conditions were: 5 min 95°C then (1 min 95°C, 1 min 58°C, 2 min 72°C) x 42 cycles. PCR products were purified from agarose gels (NucleoSpin Gel and PCR Clean-up, Macherey-Nagel). Nested PCR (position, 168303-168445) was performed with 1/100 of the purified first round PCR products, primers were 5’ CTATCATCTACTCAATGTCTATTAGACG and 5’GTTCTGTACATTGATCATATATAACTACTC, amplification conditions were: 5 min 95°C then (30 sec, 78-82°C, 30 sec 58°C, 1 min 72°C) x 45 cycles, then 20 min 72°C.

For the PCR amplification of MPXV from patients, the first-round primers were identical to the *in vitro* amplification. Nested PCR (nucleotide 168240 to 168487) was performed with 1/100 of the purified first round PCR products, primers were 5’ TAACGCCCTTGGCTCTAACCATTTTCAA and 5’GACGTGTTTGTTGAGTATCGGTGATAA, amplification conditions were: 5 min 95°C then (30 sec, 76-86°C, 30 sec 58°C, 1 min 72°C) x 45 cycles, then 20 min 72°C. The choice to use different internal primers during the 3D-PCR concerning the experiments performed *in vitro* provided a higher heterogeneity of MPXV populations, thus facilitating the mutational analysis.

3D-PCR products were purified from agarose gels (NucleoSpin Gel and PCR Clean-up, Macherey-Nagel) and ligated into the TOPO TA cloning vector (Invitrogen), and ~20-100 colonies were sequenced. A minimum threshold of two G->A transitions per sequence was imposed to reduce the impact of MPXV natural variation and PCR error in designating A3 editing.

### Phylogenetic analyses

We retrieved all complete or near-complete MPXV sequences available on GenBank as of July 4th, 2022. Sequences were aligned to reference sequence (accession MT903344) using MAFFT v7.467 (option --thread 20 --auto --keeplength --addfragments), and positions 0-1500, 2300-3600, 6400-7500, 133050-133250, 173250-173460 and 196233-end were masked (replaced with N) using goalign v0.35 (Lemoine and Gascuel, 2021) (options mask -s <start> -l <length>). From this multiple sequence alignment, a phylogenetic tree with bootstrap supports was inferred using IQ-TREE 2 v2.0.6 (options -nt 20 --safe -m GTR+G4+FO --seed 123456789 -b 100) (Minh et al., 2020). The tree was rooted using the sequence NC_003310 as outgroup (concordant with the root detected by temporal signal using Tempest v1.5.3 (Rambaut et al., 2016)). Ancestral sequences at each internal node of the phylogeny were inferred using raxml-ng v1.0.1 (Stamatakis, 2014) (options --ancestral --msa <msa> --tree <iqtree tree> --model GTR+G4+FO), and all mutations corresponding to branches of interest were counted using a dedicated script. To compute the expected number of each type of mutation, we simulated sequences along the tree (at internal nodes and tips), 100 times, using seq-gen v1.3.4 (https://github.com/rambaut/Seq-Gen), taking the inferred ancestral sequence as the root sequence, with GTR model and the parameter values (rate parameters, state frequencies and alpha parameter of the gamma distribution) initially optimized by IQ-TREE 2 while inferring the tree. We then counted the mutations between internal node sequences corresponding to branches of interest and averaged the number of mutations over the 100 simulations.

### Sanger sequencing data analysis

We preprocessed the isolate sequence (GenBank accession ON755039), to keep the sub-sequence from positions 168239 to 168487 (numbering according to the reference sequence, GenBank accession MT903344), using goalign v0.3.6a (options subseq -s <start> -l <length>). We then mapped the sequences of all the samples to this reference using minimap v2.24 (Anderson et al., 2018) (options -a -x splice --sam-hit-only --secondary=no --score-N=0). We converted the sam files to a multiple sequence alignment using gofasta v1.1.0 (Jackson, 2022) (options sam toMultiAlign -s <sam> --start 168240 --end 168487. We then removed sequences having more than 70% N using goalign v0.3.6a (options clean seqs --char N -c 0.7). Alignment and Sanger sequence traces were visually inspected using Geneious Prime (Biomatters Ltd.) We finally counted the different mutations and their context on the reference sequence using a python script (https://github.com/Simon-LoriereLab/MPXV_apo).

### Ethical statement

The IRBA is the national reference center for orthopoxviruses (CNR-LE orthopoxvirus), designated by the French Ministry of Health (through the *“Arrêté du 7 mars 2017 fixant la liste des centres nationaux de référence pour la lutte contre les maladies transmissibles”*) to process the samples for identification and characterization of MPXV. The patient signed an informed consent, and the sample subjected to viral genetic characterization was processed in an anonymized fashion. All work with infectious virus was performed in BSL-3 containment laboratories.

## Data availability

All data supporting the findings of this study are available in the manuscript.

Supplementary Information is available for this paper.

## Acknowledgments

We would like to thank all the healthcare workers, public health employees, and scientists involved in the Monkeypox outbreak response. We acknowledge the authors, originating and submitting laboratories of the MPXV sequences from GenBank. This work used the computational and storage services provided by the IT department at Institut Pasteur, Paris.

## Author contributions

RS and JPV designed research, RS, KR, SG, LB performed experiments, FL and ESL performed phylogenetic analysis, RS, KR, SG, LB, FL, OF, JNT, FI, ESL and JPV analyzed data, RS, KR, ESL and JPV wrote the paper with inputs from all authors.

## Figure legends

**Figure supplement 1**

**Phylogeny of MPXV genomes.**

The same phylogeny of MPXV genomes as shown in Figure 1, presenting all sequences and their names, as well as bootstrap supports. Branches with a dark blue circle are supported at 100%, and branches with a light blue circle are supported at ≥ 90%.

**Figure supplement 2**

**Hypermutation of MPXV DNA *in vitro* and *in vivo*.**

a) Complete G->A and C->T hypermutants resulting from A3F deamination of MPXV DNA obtained with a Td=80.7°C. b) C->T hypermutant of MPXV DNA from patient 17 with a Td=95°C. c) Complete G->A and C->T hypermutants of MPXV DNA from patient 17 with a Td=79.9°C. d) Complete G->A and C->T hypermutants of MPXV DNA from patient 21 with a Td=79.9°C. the “Gs” highlighted in bold red in the reference sequences represent the deamination hotspot sites described in Figure 3b. The sequence is given with respect to the viral plus strand. Only differences are shown. Hyphens denote gaps introduced to maximize sequence identity. All sequences were unique, indicating that they corresponded to distinct molecular events. The dots in the sequences represent identical bases to the reference sequence. the values on the left and right of the sequences represent respectively the sequence number and the amount of mutations per sequence.

**Figure 2-source data 1**

Raw images of the 3DPCR agarose gel in Figure 2a. The dashed boxes indicate the area of gels presented in the Figure.

**Figure 3-source data 1**

Raw images of the 3DPCR agarose gel in Figure 3c. The dashed boxes indicate the area of gels presented in the Figure.

**Table supplement 1**

Presentation of the 24 studied patients, specifying gender, date of birth, type of sampling, type of lesions and anatomical localization of analyzed lesions.

